# Cell surface RNA expression modulates alveolar epithelial function

**DOI:** 10.1101/2024.05.19.594844

**Authors:** Jubilant Kwame Abledu, Christopher J. Herbst, Raphael Brandt, Alen Kocak, Pritam Ghosh, Jacob L. Gorenflos López, Kevin Diestelhorst, Stephan Block, Christian Hackenberger, Oliver Seitz, Elena Lopez-Rodriguez, Matthias Ochs, Wolfgang M. Kuebler

## Abstract

Glycosylated RNA (glycoRNA) has recently emerged as a novel constituent of the glycocalyx on cell surfaces, yet its biological functions remain largely unexplored. In this report, we present the first analysis of glycoRNA expression and functionality in alveolar epithelial cells.

To this end, we optimized new techniques for the detection of glycoRNA on living cell surfaces and in cell membrane-associated RNA samples through in-gel imaging after labeling with fluorescent dye conjugates. Specifically, we used conjugation of Cy5-hydrazide following mild oxidation with sodium periodate for detection of the entire cell surface sialoglycoRNA pool. Conjugation of dibenzocyclooctyne-sulfo-Cy5 (DBCO-Sulfo-Cy5) in cells fed with tetraacetylated *N*-azidoacetyl-mannosamine (Ac_4_ManNAz) or 6-azido-L-fucose (FucAz) detected *de novo* formed sialoglycoRNA or fucoglycoRNA. Finally, biotinylated lectins in combination with infrared dye-conjugated streptavidin was used to differentiate between specific glycosidic linkages.

Comparisons across primary alveolar epithelial cells and different alveolar-epithelial like cell lines revealed a cell-type specific variation in glycoRNA abundance. Treatment of primary alveolar epithelial cells with an RNAse cocktail reduced epithelial surface glycoRNA and was associated with a reduction in trans-epithelial electrical resistance and influenza A viral particle abundance.

As such, the present work identifies glycoRNA as a novel component of the alveolar epithelial glycocalyx, suggesting its potential relevance in epithelial barrier regulation and viral infection.

## Introduction

The alveolar epithelium plays a pivotal role in facilitating lung gas exchange while concurrently serving as a physiological barrier to prevent the uncontrolled leakage of fluid and proteins into the air space and restricting the entry of inhaled pathogens into the bloodstream. It is composed of two anatomically and functionally distinct cell types: i) squamous alveolar epithelial type I cells (AEI) which cover about 95% of the alveolar epithelial surface and ensure efficient gas exchange, and ii) cuboidal alveolar epithelial type II cells (AEII) which synthesize and secrete surfactant and serve as progenitor cells for AEI cells. On its apical side, the alveolar epithelium is coated with a glycocalyx – a carbohydrate-rich cell surface coat consisting of glycoproteins, glycolipids, glycosaminoglycans and proteoglycans ^1–3^.

Previous work has identified important roles of the glycocalyx as regulator of cell and organ function including barrier stability, mechanosensation, infection and immunity ^4,5^. The majority of these effects have been demonstrated in the vascular endothelium, while the functions of the alveolar epithelial glycocalyx are less clear. E.g. degradation of glycosaminoglycans (GAGs) of the alveolar epithelial glycocalyx has been shown to increase alveolar protein permeability yet without causing lung edema ^1^, presenting a seeming paradox. As such, the composition of the alveolar epithelial glycocalyx and the individual effects of its constituents on alveolar epithelial function require a more detailed understanding.

Recently, Flynn and colleagues identified glycosylated RNA as a novel component of the glycocalyx in living cells and organs ^6^. These so-called glycoRNAs comprise small non-coding RNAs that are expressed on the cell surface and modified by sialic acid- and fucose-containing *N*-glycans. The authors further demonstrated that glycoRNAs interact with immunomodulatory receptors, specifically the sialic acid binding-immunoglobulin-like lectins (Siglecs) 11 and 14, suggesting a potential role of glycoRNAs as adhesion receptors for neutrophils, monocytes, and macrophages. Likewise, immune cells have been shown to similarly express glycoRNA ^6,7^ which in turn modulates their interaction with other cells recently demonstrated for the interaction between monocytes/macrophages and vascular endothelial cells ^7^. Yet, the expression and functional role of glycoRNA as constituent of the alveolar epithelial glycocalyx remains so far unexplored.

Here, we provide the first analysis of the expression of glycan-associated RNA on the surface of alveolar epithelial cells, its composition in terms of glycan residues and glycosidic bonds, and its implication on epithelial function. While we will refer to these entities as glycoRNAs from hereon in analogy to previous work, the present study did not focus on the molecular structure of these glycan-associated RNAs, but primarily on their detection and function in the alveolar epithelium. To this end, we have optimized a set of techniques for the rapid labeling and visualization of cell surface glycoRNAs in denaturing agarose gels using fluorescent probes. Specifically, we have focused analyses on cell surface expressed rather than bulk glycoRNAs, avoided signal loss by labeling glycoRNAs with fluorescent probes and subsequent in-gel imaging, and developed different approaches to probe for both overall expressed or newly formed cell surface glycoRNAs, as well as for different glycan moieties and glycosidic bonds. To this end, we first labeled *de novo* surface-expressed glycoRNA with dibenzocyclooctyne-sulfo-Cy5 (DBCO-Sulfo-Cy5) using click chemistry subsequent to feeding cells with the non-natural azide analogues of the sialic acid or fucose precursors *N*-acetyl mannosamine (ManNAc) and L-fucose, namely tetraacetylated *N*-azidoacetyl-mannosamine (Ac_4_ManNAz) and 6-azido-L-fucose (FucAz), respectively. Sialylated total cell surface glycoRNA was labeled with Cyanine5-hydrazide (Cy5-hydrazide) following mild oxidation with sodium periodate. Finally, glycan residues and glycosidic bonds were identified by cell surface glycoRNA labeling with biotin-labeled lectins that were detected by infrared dye-conjugated streptavidin.

Testing of these methods across commercially available primary alveolar epithelial cells or alveolar-epithelial like cell lines, respectively, revealed a cell-type specific variation in glycoRNA abundance. Finally, treatment of primary alveolar epithelial cells with an RNAse cocktail revealed important functional roles of glycoRNAs, in that RNAse treatment reduced both epithelial barrier function and the adhesion of virus particles to the cell surface. As such, the present work identifies glycoRNAs as a novel constituent of the alveolar epithelial glycocalyx with important functional relevance for epithelial barrier regulation, infection and immunity.

## Methods

### Cell culture and chemicals

Cells were grown at 37° and 5% CO_2_ in T25 or T75 cell culture flasks (Falcon), 35 mm petri dishes (Greiner Bio-One), or 6-well cell culture plates (Falcon) where necessary. Human primary alveolar epithelial cells (hPAEpC, Cell Biologics) at passages 4 to 7 were cultured in complete human alveolar epithelial medium (Cell Biologics). Human alveolar epithelial lentiviral immortalized cells (CI-hAELVi, InSCREENeX GmbH) were cultured in complete human alveolar epithelial cell medium (InSCREENeX GmbH) at passages 13 to 35. The lung adenocarcinoma-derived cell lines, A549 (passages 17 to 28) and H441(passages 16 to 30), were respectively cultured in Gibco DMEM and RPMI mediums supplemented with 10% fetal bovine serum (FBS) and 1% penicillin/streptomycin (P/S).

Stocks of 100 mM Ac_4_ManNAz (Jena Bioscience), 200 mM 6-Azido-L-Fucose (Synthose), 100 mM dibenzocyclooctyne-polyethylene-glycol-4-biotin (DBCO-PEG_4_-biotin) (Jena Bioscience) and 25 mM DBCO-Sulfo-Cy5, DBCO-AF555 and DBCO ATTO-488 (all Jena Bioscience) were generated in sterile dimethyl sulfoxide (DMSO).

Stocks of glycan biosynthesis inhibitors were generated in DMSO at the following concentrations: 10 mM NGI-1 (Tocris), 10 mM kifunensine (Sigma), 100 mM P3-F_AX_-Neu5Ac (Tocris). All compounds were stored at -20^°^ until use. For periodate oxidation and aldehyde labeling, the following stocks were generated: NaIO_4_ 100 mM (ThermoFisher Scientific) in deionized water, glycerol 100 mM (ThermoFisher Scientific) in deionized water, 2-amino-5-methoxybenzoic acid 1 M (ThermoFisher Scientific) in DMSO, Cy5-hydrazide 25 mM (Lumiprobe) in DMSO.

Stock of 1 mg/ml Calcein AM (ThermoFisher Scientific) was generated in sterile DMSO.

### Metabolic labeling of glycoRNA

Cells were fed with either 100 μM Ac_4_ManNAz or 200 μM FucAz for 24 h. Cells were washed twice with 1xPBS and then incubated with 100 μM DBCO-PEG_4_-biotin, allowing for the detection of newly formed sialic acids or fucose residues on the cell surface by click chemistry ^8^.

### Detection of glycoRNA by in vitro click conjugation

GlycoRNA was labelled and detected *in vitro* as described previously with a few modifications ^6^. Following feeding with Ac_4_ManNAz or FucAz for 24 h, cells were washed twice with 1xPBS and lysed by RNAzolRT (Molecular Research Inc). RNA was extracted and purified with Zymo RNA clean and concentrator kit (Zymo Research). Thirty micrograms of RNA in 9 μL pure water was mixed with 10 μL of RNA denaturing buffer (95% formamide, 18 mM EDTA and 0.025% SDS) and 1 μL of DBCO-PEG_4_-biotin (10 mM; Jena bioscience). Conjugation of DBCO to azide residues was performed at 55°C for 10 min and the reaction was stopped by adding 80 μL water, followed by 200 μL RNA binding buffer (Zymo Research), vortexing, and finally addition of 300 μL isopropanol and vortexing. The final mixture was purified with Zymo RNA clean and concentrator kit and used for RNA extraction as described below.

### Enzymatic digestions

All enzymatic digestions were performed on 30 μg of RNA in 50 μL at 37°C for 1 h. To digest RNA, 1 μL of RNAse cocktail (RNAse A= 500 U/mL, RNAse T1= 20,000 U/mL, ThermoFisher Scientific) was used in 20 mM Tris-HCl (pH 8.0), 100 mM KCl and 0.1 mM MgCl_2_. Glycan digestion was achieved with the following enzyme concentrations: 5 μL of α2-3,6,8,9 neuraminidase A (20 U/μL, New England Biolab, NEB) with 1x GlycoBuffer 1 (NEB), 5 μL of PNGaseF (500 U/μL, NEB) with 1x GlycoBuffer 2 (NEB), 5 μL of O-glycosidase (40,000 U/μL, NEB) with 1x GlycoBuffer 3 (NEB), 5 μL of α1-2,4,6 fucosidase O (2U/μl, NEB) with 1x GlycoBuffer 1 (NEB).

### Periodate oxidation and aldehyde labeling (pAL) of glycoRNA

As an alternative strategy for glycoRNA labeling, we targeted cell surface sialic acids as recently described, with a few modifications ^9^. Cells were washed twice with 1x PBS and then subjected to mild periodate oxidation by adding 1 mM NaIO_4_ (ThermoFisher Scientific) for 15 minutes at room temperature. The periodate was removed and quenched by adding 1 mM glycerol (ThermoFisher Scientific) for 1 minute, followed by two additional washes with 1x PBS. Next, 25 μM Cy5-hydrazide with 10 mM 2-amino-5-methoxybenzoic acid (ThermoFisher Scientific) in 1xPBS as a catalyst was added to the cells for 15 minutes at room temperature to allow complete conjugation with the aldehyde. RNA was extracted and labeled glycoRNA detected by electrophoresis and in-gel imaging as described below.

### Plasma membrane isolation

Plasma membrane was isolated according to previously published protocols.^10^ After culturing to confluence in T175 cell culture flasks (Falcon), cells were scraped into microtubes containing cold homogenization buffer (0.25 M sucrose, 20 mM HEPES-KOH, pH 7.4, and a protease inhibitor mixture). The cell suspension was sonicated on ice using alternating on and off pulses of 5 and 10 seconds for approximately 30 rounds. Subsequently, the sample was centrifuged at 1,000 x g for 10 minutes to remove the nuclear fraction and debris. The resulting supernatant was subjected to two rounds of ultracentrifugation at 200,000 x g for 45 minutes each, with a washing step using Na_2_CO_3_ (pH 11) after the first round. The obtained membrane pellets were homogenized in 200 μL of RNAzol (Molecular Research Inc) and processed for RNA extraction (*vide infra*).

### GlycoRNA labeling with lectins

To identify specific glycosidic bonds, lectin labeling was performed on 30 μg of plasma membrane-derived RNA in 500 μL at 37°C for 20 min. RNA was combined with 20 μg/ml biotinylated lectin – wheat germ agglutinin (WGA, EY laboratory), *Maackia amurensis* agglutinin (MAA, EY laboratory), *Sambucus nigra* agglutinin (SNA, EY laboratory) or *Ulex europaeus* agglutinin 1 (UEA 1, EY laboratory) – and 1 μL of infrared dye-conjugated streptavidin (IRDye Streptavidin 800CW, LI-COR Biosciences) with 1x GlycoBuffer 1 (NEB). Following incubation, the mixture was combined with 1x RNA binding buffer (Zymo Research), 2 x isopropanol and purified with Zymo RNA clean and concentrator kit (Zymo Research) for subsequent RNA purification.

#### RNA extraction and purification

RNA was extracted with RNAzol RT (Molecular Research Inc) as previously described ^11^. In brief, 1 ml of RNAzol was added to the cell culture dish/well to lyse cells and the cell lysate was homogenized by pipetting. Phase separation was performed by adding 0.4 ml of RNAse-free water, vortexing, incubating at room temperature for 5 min, and finally centrifugation at 12,000 x g at 4°C for 15 min. The RNA-containing aqueous phase was carefully transferred into a clean tube, mixed with 1.1x volume of isopropanol and purified over Zymo columns (Zymo Research). First, the column was preconditioned by adding 350 μL of pure water, spun at 10, 000 x g for 20 seconds, and the flowthrough was discarded. Next the precipitated RNA was spun through the column at 10, 000x g and the flowthrough was again discarded. Three separate washes were performed on the column: one time with 400 μL RNA prep buffer (3 M guanidine hydrochloride in 80% ethanol) and two times with 80% ethanol, followed by spinning at 10,000 x g for 20 seconds (for the first two washes) and 30 seconds (for the last wash), respectively. To digest proteins, 1.05 μg/μL Proteinase K (ThermoFisher Scientific) was diluted in water and added directly to the column (50 μL for Zymo-II and 60 μL for Zymo-IICG). The column was sealed with a cap or parafilm and then incubated at 37°C for 45 min. After protein digestion, the column was brought to room temperature for 5 min and the RNA was spun out into fresh tubes (50 μL for Zymo-II and 60 μL for Zymo-IICG columns). Next, the extracted RNA was subjected to mucinase digestion by adding 1.5 μg of mucinase to 50 μL of RNA solution and incubation at 37°C. The mixture was combined with 1x RNA binding buffer (Zymo Research), 2x isopropanol and purified with Zymo RNA clean and concentrator kit (Zymo Research). The purified RNA was used for electrophoresis as described in the next paragraph.

#### GlycoRNA blotting

GlycoRNA blotting was performed with few modifications according to the previously published protocol by Flynn and colleagues ^6^. In brief, DBCO-PEG_4_-biotin-labelled RNA was lyophilized dry, resuspended in 15 μL of Gel Loading Buffer II (95% formamide, 18 mM EDTA, 0.025% SDS) with 1x SybrGold (Thermo fisher), incubated at 55^°^C for 10 min and finally placed on ice for 3 min. Three micrograms of RNA was loaded into 1% agarose-formaldehyde gel and electrophoresed in 1x MOPS buffer at 115V for 35 minutes. Total RNA was visualized on ChemiDoc MP Imaging Systems (Biorad). RNA was transferred onto 0.45 μM nitrocellulose (NC) membranes as per the Northern Max protocol for 10 h at 25°C. After transfer, the NC membrane was rinsed in 1x PBS and dried on Whatman paper. RNA crosslinking was achieved by heating the NC membrane in an oven at 80°C for 15 minutes. The NC membrane was rehydrated by rinsing in 1x PBS, blocked with Intercept PBS blocking buffer (Li-COR Biosciences) at 25°C for 45 minutes and stained with IRDye streptavidin 800CW (Li-COR Biosciences) in blocking buffer (1:10,000) for 45 minutes at 25°C. The NC membrane was washed three times with 0.1% Tween-20 in 1x PBS and then once with 1x PBS. The NC membrane was scanned on a ChemiDoc MP Imaging Systems (Biorad) for visualization of glycoRNA bands.

#### In-gel imaging of glycoRNA

Three micrograms of DBCO-Sulfo-Cy5-, Cy5-hydrazide-, or lectin-labelled RNA was lyophilized dry and combined with 9 μL of Gel Loading Buffer II (95% formamide, 18 mM EDTA, 0.025% SDS) with a 1x SybrGold (Thermo Fisher), incubated at 55°C for 10 min and finally cooled on ice for 2 min. RNA was loaded into 1% agarose, 0.75% formaldehyde, 1.5% MOPS denaturing gel and electrophoresed in 1% MOPS at 115 V for 35 minutes. The gel was scanned on ChemiDoc MP Imaging Systems (Biorad) to visualize RNA (SybrGold channel) and glycans (Cy5 channel or Streptavidin infrared).

#### Epithelial barrier function

Barrier function was assessed in hPAEpC cultures by trans-epithelial electrical resistance (TEER, EVOM2, World Precision Instruments, UK) in the presence of 50 μL/mL RNAse cocktail (ThermoFisher Scientific) or Hank’s balanced salt solution (gibco 1xHBSS, ThermoFisher Scientific) for 30 min at 37°C, respectively, as recently described ^12^.

#### Influenza virus interaction

Influenza A (IAV) strain A/WSN/1933 was produced as described previously ^13^. UV-inactivated IAVs were labeled by octadecyl rhodamine B chloride (R18; ThermoFisher Scientific) as follows: Five μL of influenza A/WSN/1933 (total protein content of 0.51 mg/mL) were diluted in 95 μL of 1x PBS buffer, followed by addition of 2 μL R18 solution (0.2 mM in ethanol) and gentle mixing. The mixture was stored on ice for 30 min in the dark. Free dye (i.e., R18 molecules, which did not incorporate in the IAV envelope) was removed using a microspin column PD SpinTrap G-25 (Cytiva) at 3300 rpm for 2 min.

To probe for a possible interaction between viral particles and glycoRNA, we treated confluent hPAEpC in 18-well chamber slides (μ-Slide 18 well, Ibidi) with 50 μL/mL RNAse cocktail (ThermoFisher Scientific) or HBSS for 30 min at 37°C. After two washes with HBSS, cells were incubated with 2 μg/mL R18-labelled IAV in culture medium for 1 h at 37°C. Cells were then washed twice with HBSS, fixed in 4% PFA for 20 minutes, and viral particles interacting with hPAEpC were visualized by spinning disk confocal microscopy (Nikon Instruments Inc.).

#### Cell viability

To evaluate the impact of RNAse on cell viability, hPAEpC were cultured to confluence in a 24-well plate (Greiner Bio-One). After treatment with RNAse cocktail for 30 minutes at 37°C, cells were stained with calcein AM (1:1,000, ThermoFisher Scientific) and propidium iodide (1:1,000, ThermoFisher Scientific) in culture media for 30 minutes at room temperature. The proportion of dead cells (stained with propidium iodide) and live cells (stained with calcein AM) was examined using a fluorescent plate imager (ZOE).

#### Statistical analysis

Statistical analyses were performed using GraphPad Prism 10 (GraphPad Software Inc., USA). Statistical tests used and n-number of replicates are specified in each figure legend; differences with p<0.05 were considered significant.

## Results

### GlycoRNA is expressed on the cell surface of alveolar epithelial cells

Conventional approaches for labeling glycoRNA in denaturing agarose gels entail several sequential steps, including bulk RNA extraction, enzymatic purification, labeling, and electrophoresis ^6^. Yet, as these labeling techniques cannot differentiate between cytosolic and cell surface glycoRNA, we designed a novel protocol to directly label cell surface glycoRNA *in situ* before RNA extraction and electrophoresis. To this end, hPAEpC were first subjected to metabolic labeling with Ac_4_ManNAz or FucAz, which can enter the sialic acid or fucose biosynthesis pathway, respectively, resulting in incorporation of azide-modified sialic acids or fucose in *de novo* formed glycoRNA. Subsequently, cells were treated with the click-chemistry compound, DBCO-PEG_4_-biotin for biotinylation of sialic acid- and fucose-containing glycoRNA as illustrated in Figure 1A. The extracted and purified RNA was subjected to denaturing gel electrophoresis and blotting, and the biotin-labeled glycoRNA was detected by streptavidin dye. Sialylated glycoRNA (denoted as sialoglycoRNA in the following) was not only detected in conventionally extracted bulk RNA from whole hPAEpC, but also by specific cell surface labeling as described above (Figure 1B), while signals were absent in cells not fed with Ac_4_ManNAz. Surface expression of *de novo* formed sialoglycoRNA increased as a function of time over the duration of Ac_4_ManNAz treatment (Figure 1C) but was reduced in the presence of the sialyltransferase inhibitor P3-F_AX_-Neu5Ac (Figure 1 D). The sialoglycoRNA signal was abolished by treatment of purified RNA with RNAse or neuraminidase (Figure 1E). Similarly, sialoglycoRNA was no longer detected when purified RNA had been treated with the *N*-glycan-specific glycosidase, peptide: N-glycosidase F (PNGaseF), yet was not affected by treatment with *O*-glycosidase, which specifically cleaves serine- or threonine-linked *O*-glycosides (Figure 1F). Consistent with these findings, sialoglycoRNA expression on the surface of alveolar epithelial cells was sensitive to treatment with chemical inhibitors of *N*-glycosylation, such as NGI-1 (Figure 1G) and kifunensine (Figure 1H), indicating that sialoglycoRNAs primarily contain similar bonds as *N*-linked glycans. In parallel, we observed a time-dependent expression of fucosylated glycoRNA (or fucoglycoRNA) on the epithelial cell surface following FucAz treatment (Figure 1I) that was again abolished by RNAse, fucosidase or PNGaseF but not by *O*-glycosidase treatments (Figure 1J). FucoglycoRNA was again sensitive to NGI-1 and kifunensine treatment. Collectively, our data reveal the expression of both sialoglycoRNA and fucoglycoRNA on the cell surface of alveolar epithelial cells.

**Figure 1:**
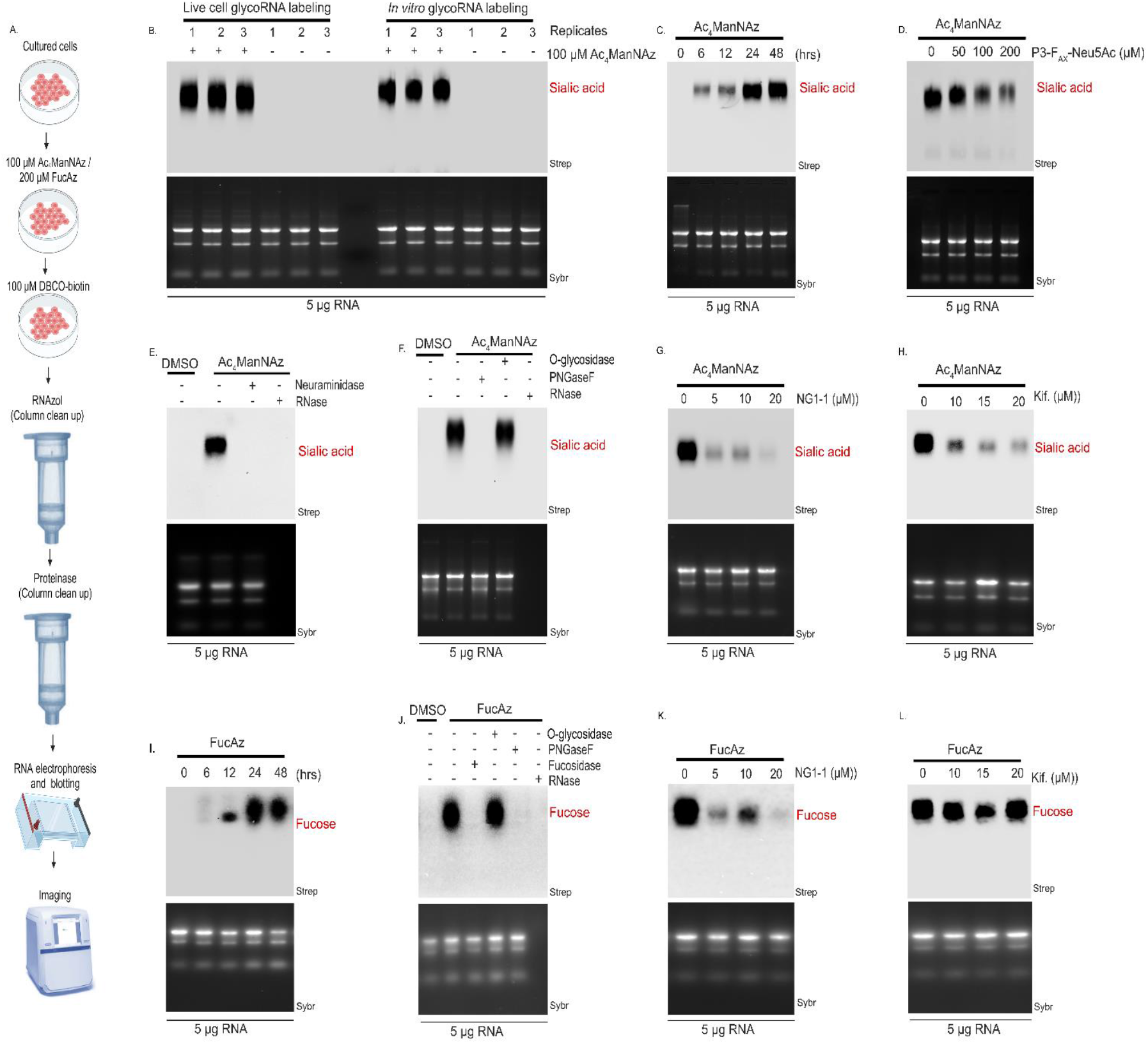
Metabolic labeling with Ac_4_ManNAz and FucAz reveals the expression of cell surface glycoRNA in human primary alveolar epithelial cells. (A) Schematic of RNA labeling with Ac_4_ManNAz and FucAz in human primary alveolar epithelial cells, and subsequent extraction and blotting. Cells were pretreated with 100 μM Ac_4_ManNAz or 200 μM FucAz for 24 h followed by 25 μM DBCO-PEG_4_-biotin. After purification, 5 μg RNA was loaded onto 1% denaturing agarose gels, subjected to electrophoresis for 35 min, and imaged on a fluorecscent gel scanner. In control experiments, glycosidases were replaced with RNase-free water while chemical inhibitors were replaced with DMSO. (B) Blotting of RNA from hPAEpC treated with 100 μAM Ac_4_ManNAz for 24 h. Following RNA purification, DBCO-biotin was added to either live hPAEpC (left; for cell surface glycoRNA labeling) or bulk RNA from hPAEpC (right, for *in vitro* labeling of the total glycoRNA pool). RNA was transferred onto nitrocellulose membranes, and sialoglycoRNA was visualized with streptavidin-IR800 CW (Strep) on a fluorescent scanner. RNA was stained and imaged with SYBR Gold (Sybr). (C) Blotting as in (B) for cell surface glycoRNA in hPAEpC treated with 100 μM Ac_4_ManNAz for indicated times shows *de novo* formation of sialoglycoRNA. (D) Blotting in hPAEpC treated with 100 μAM Ac_4_ManNAz in the presence of the sialyltransferase inhibitor P3-F_Ax_-Neu5Ac shows dose-dependent inhibition of glycoRNA sialylation. (E) Blotting as in (B) in the presence of neuraminidase (2 U/μL) or RNAse cocktail (1:50). (F) Blotting as in (B) in the presence of *O*-glycosidase (4000 U/μL), the N-glycan specific glycosidase, *PNGAseF* (50 U/μL) or RNAse. (G) Blotting as in (B) in the presence of oligosaccharyltransferase (OST) inhibitor NGI-1 at the given concentrations. (H) Blotting as in (B) in the presence of kifunensine (Kif), a chemical inhibitor of the N-glycan trimming enzyme α-amannosidase I, at the given concentrations. Blots (D-H) were obtained in cells fed with 100 μM Ac_4_ManNAz for 24h. (I) Blotting as in (C) in hPAEpC treated with 200 μAM FucAz for indicated times shows *de novo* formation of fucoglycoRNA. (J) Blotting as in (I) in the presence of *O*-glycosidase, *PNGAseF*, fucosidase (0.2U/μl) or RNAse. (K) Blotting as in (I) in the presence NGI-1 at the given concentrations. (L) Blotting as in (I) in the presence of the kifunensine (Kif) at the given concentrations. Blots (J-L) were obtained in cells fed with 200 μM FucAz for 24h. Created with Biorender.com

### Click chemistry labeling with DBCO-Sulfo-Cy5 enables rapid in-gel detection of *de novo* formed glycoRNA

A persistent challenge associated with conventional RNA blotting techniques is the incomplete transfer of RNA onto nitrocellulose membranes. Although recent studies proposed enhanced RNA transfer efficiency by use of optimized transfer buffers (3 M NaCl, pH<2 or >12), ^11^ we only achieved complete RNA transfer after extended incubation periods of 10-18 hours, while transfer within the previously specified 2.5-hour period was incomplete (data not shown). To circumvent this problem, we considered a labeling approach using fluorescently-conjugated DBCO instead of biotin that would facilitate the direct visualization of glycoRNA within the gel, thereby eliminating the need for RNA transfer altogether. Specifically, we considered that DBCO-conjugated fluorescent dyes with excitation (Ex)/emission (Em) wavelengths within or beyond the far-red spectral region (i.e., ≥ 600/650 nm) such as Cy5 (Ex=649 nm, Em=667 nm) would enable the simultaneous visualization of glycans (i.e., sialic acids or fucose, stained with Cy5) and RNA (stained with SYBR Gold with Ex=496 nm, Em=539 nm) without spectral overlaps. Accordingly, we labeled glycoRNA with DBCO-Sulfo-Cy5 after feeding the cells with Ac_4_ManNAz or FucAz as illustrated in Figure 2 A. RNA was extracted and purified as described before; following denaturing gel electrophoresis glycoRNA was visualized directly in-gel. DBCO-Sulfo-Cy5 yielded a robust and stable (i.e. without obvious bleaching over time) glycoRNA labeling. Pilot experiments with co-staining of glycoRNA by DBCO-Alexa Fluor 555 or DBCO-ATTO 488 for glycans and SYBR-Gold for RNA showed considerable spectral overlap (with RNA signals conspicuously appearing in the glycan spectrum) with very weak glycan signals, yet this problem could be prevented by the combination of Cy5 (for glycans) and SYBR-Gold (for RNA) (Figure 2B). As would be expected from our previous results, sialoglycoRNA labeled with DBCO-Sulfo-Cy5 was again effectively removed by treatment of purified RNA with neuraminidase, RNAse (Figure 2C) or PNGaseF, while treatment with *O*-glycosidase had no effect (Figure 2D). On the surface of the epithelial cells, the DBCO-Sulfo-Cy5 signal was again sensitive to P-3F_AX_-Neu5Ac, kifunensine, or NGI-1 treatments, further validating its specificity for sialic acid residues (Figure 2E). Analogously, fucoglycoRNA as detected by DBCO-Sulfo-Cy5 was eliminated upon treatment with fucosidase and PNGase F, but not by *O*-glycosidase (Figure 2F). These findings validate in-gel labeling by click chemistry as a sensitive and reproducible approach for glycoRNA detection, and confirm the parallel expression of sialic acid and fucose moieties on cell surface glycoRNAs of primary alveolar epithelial cells.

**Figure 2.**
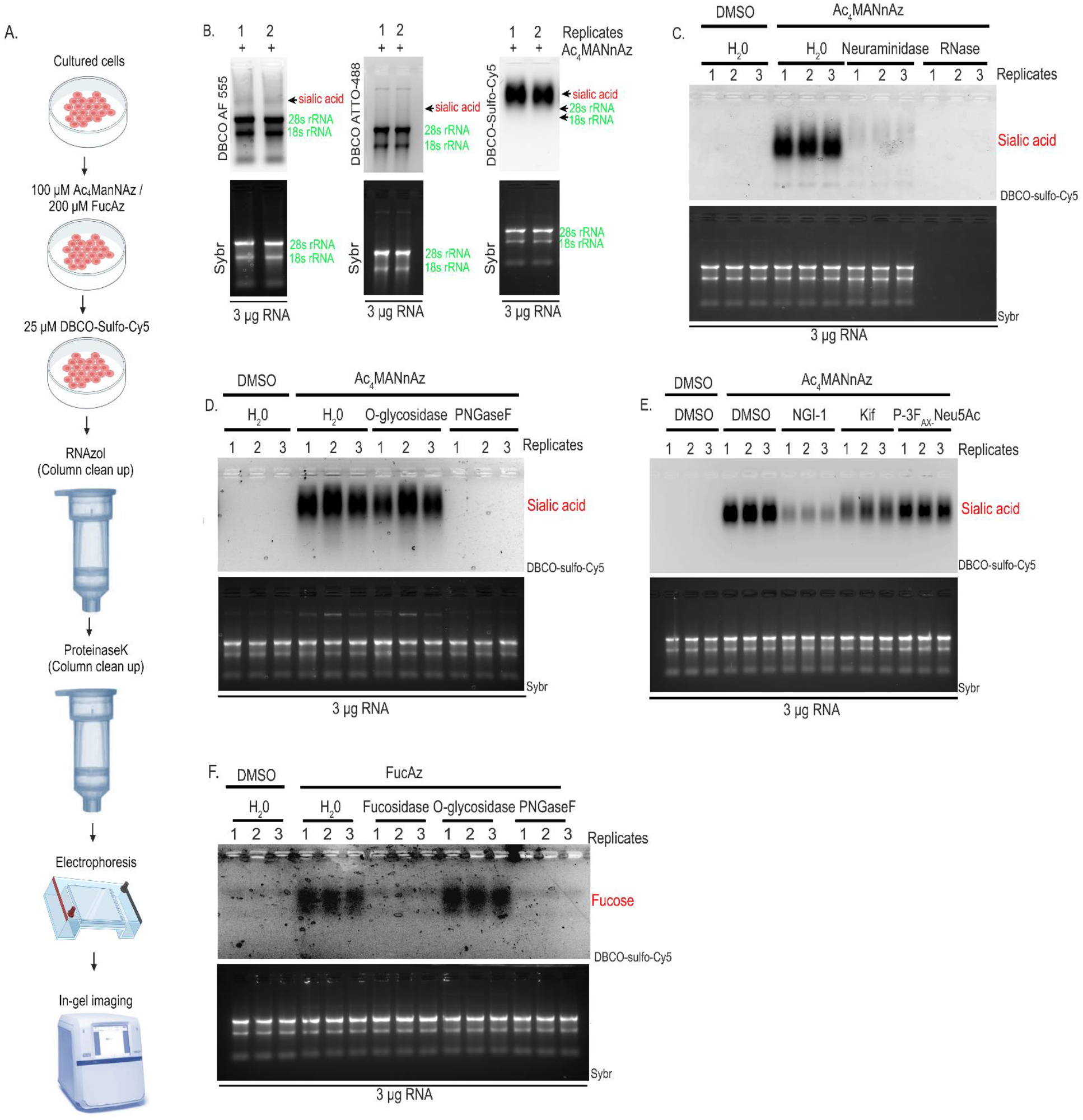
Click-chemistry labeling with DBCO-Sulfo-Cy5 enables rapid in-gel detection of *de novo* formed glycoRNA. (A) Schematic of cell surface glycoRNA labeling with Cy5-hydrazide. hpAEpC were pretreated with 100 μM Ac_4_ManNAz for 24 h followed by 25 μM DBCO-Sulfo-Cy5. After purification, 3 μg RNA was loaded onto 1% denaturing agarose gels, subjected to electrophoresis for 35 min, and imaged on a fluorecscent gel scanner. In control experiments, RNA was treated with RNAse-free water instead of glycosidases while cells were treated with DMSO instead of inhibitors. RNA was stained with SYBR gold (Sybr). (B) Comparison of different DBCO-dye conjugates for in-gel detection of sialoglycoRNA in hPAEpC show clear and distinct staining with DBCO-Sulfo-Cy5 but not with DBCO AF 555 or DBCO ATTO-488. (C-D) DBCO-Sulfo-Cy5 labeled sialoglycoRNA signal is lost in hPAEpC-derived RNA in the presence of neuraminidase (2 U/μL), RNAse (1:50) and *PNGaseF* (50 U/μL) but not *O*-glycosidase (4000 U/μL). (E) DBCO-Sulfo-Cy5 labeled sialoglycoRNA signal is reduced in hPAEpC in the presence of the oligosaccharyltransferase inhibitor NGI-1 (20 μM), the α-mannosidase I inhibitor kifunensine (Kif) (15 μM), or the sialyltransferase inhibitor P3-F_Ax_-Neu5Ac (200 μM). (F) In-gel detection of fucoglycoRNA with DBCO-Sulfo-Cy5 in FucAz-fed hPAEpC shows reduced signal in the presence of fucosidase (0.2U/μl) or *PNGaseF* (50 U/μL) but not *O*-glycosidase (4000 U/μL). Created with Biorender.com

### Periodate oxidation and Cy5-hydrazide labeling facilitate detection of the cell surface sialoglycoRNA pool

Metabolic labeling with Ac_4_ManNAz primarily targets newly synthesized sialoglycoRNA. To label the entire pool of sialoglycoRNA, sialic acids may be converted to aldehydes via mild periodate oxidation, followed by aniline-catalyzed ligation with aminooxybiotin, RNA extraction and blotting ^11^. This type of oxime ligation (i.e. the reaction of an aldehyde with an aminooxy-containing compound) is relatively stable, yet has so far only been used for bulk RNA and reaction kinetics are typically slow at physiological pH ^14^. To directly label the entire pool of cell surface sialic acids on glycoRNA under physiologically relevant conditions, we adapted a recently reported hydrazone ligation technique (i.e. the reaction of an aldehyde with a hydrazide-containing compound) ^9^. To this end, we employed a fluorescently-conjugated hydrazide (Cy5-hydrazide), which facilitates the direct visualization of sialoglycoRNA in-gel without the need for RNA transfer, as illustrated in Figure 3 A. The entirety of the labeling procedures can be completed within a span of 30 minutes, marking a substantial eight-fold reduction in the time needed for glycoRNA labeling compared to the previously proposed oxime ligation method 11. In-gel imaging of Cy5-hydrazide labeled glycoRNA yielded a clear signal with strong contrast between lanes in the absence of obvious bleaching effects or non-specific RNA labeling as previously described for oxime ligation ^11^. To validate that the Cy5-hydrazide signal indeed reflects sialoglycoRNA, we tested for its sensitivity to chemical inhibitors and degradation enzymes targeting sialylation and *N*-glycosylation as described above. Treatment of purified RNA with neuraminidase (Figure 3B), PNGaseF and RNAse effectively eliminated the Cy5-hydrazide signal, while treatment with *O*-glycosidase had no impact (Figure 3 C). Likewise, treatment of cells with P-3F_AX_-Neu5Ac, kifunensine, or NGI-1 resulted in a notable reduction in the sialoglycoRNA signal (Figure 3D). Collectively, these findings describe a novel approach for rapid detection of the entire pool of cell surface sialoglycoRNAs, and further corroborate the expression of cell surface sialoglycoRNA in human primary alveolar epithelial cells.

**Figure 3.**
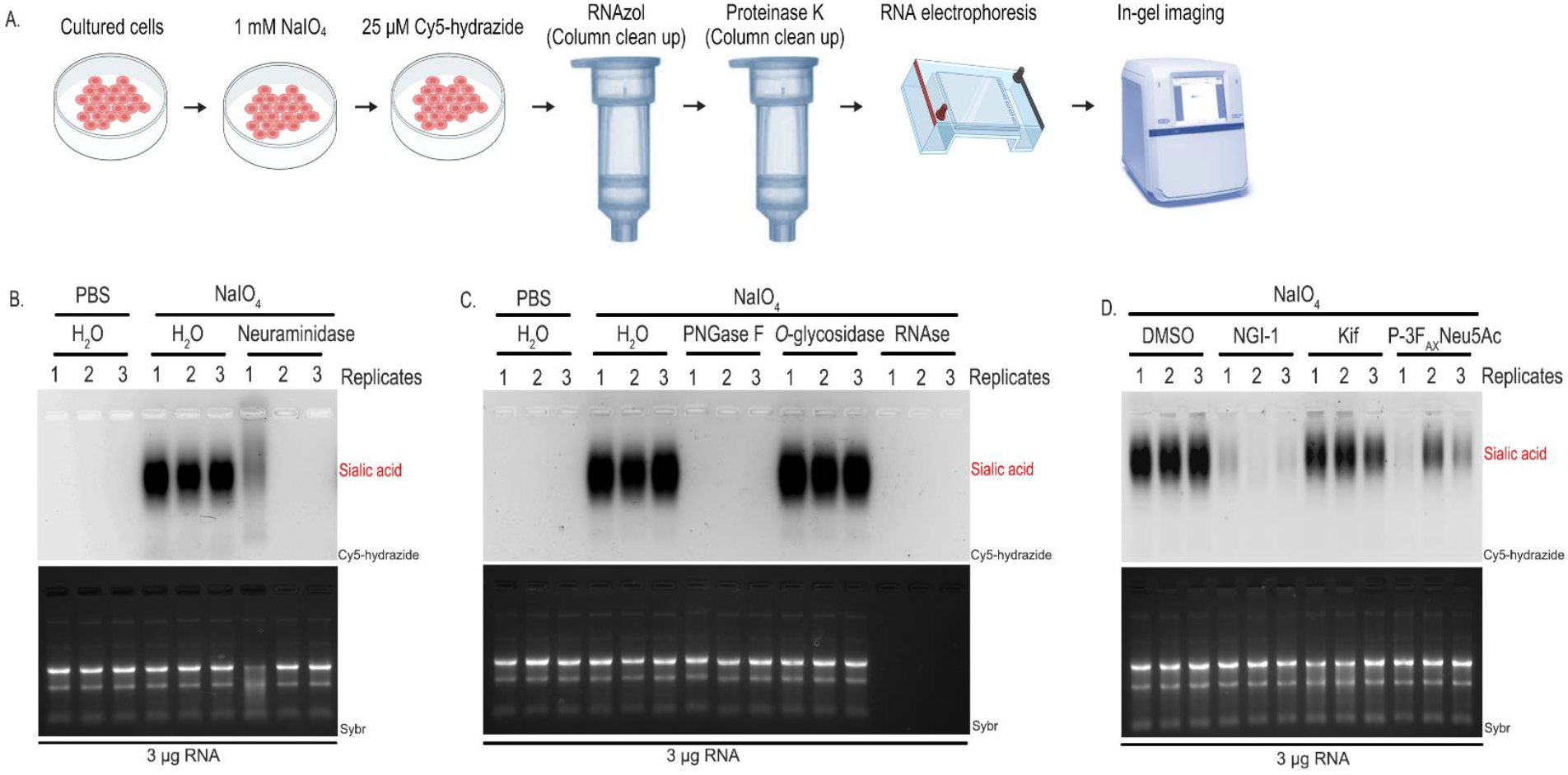
Periodate oxidation and Cy5-hydrazide labeling facilitate detection of the cell surface sialoglycoRNA pool. (A) Schematic of cell surface glycoRNA labeling with Cy5-hydrazide. hPAEpC were subjected to mild periodate oxidation with 1 mM NaIO_4_ for 15 minutes, and the reaction was quenched by 1mM glycerol followed by the addition of 25 μM Cy5-hydrazide for 15 minutes. After purification, 3 μg RNA was loaded onto 1% denaturing agarose gel, electrophoresed for 35 minutes, and imaged on a fluorescent gel scanner. All subsequent gels were treated in this manner. In control experiments, cells were treated with PBS instead of NaIO_4_ or with DMSO instead of inhibitors while RNA was treated with RNAse-free water instead of glycosidases. RNA was stained with SYBR gold (Sybr). (B) Cy5-hydrazide labeled sialoglycoRNA signal is lost in hPAEpC-derived RNA in the presence of neuraminidase (2 U/μL). (C) Cy5-hydrazide labeled sialoglycoRNA signal is lost in hPAEpC-derived RNA in the presence of *PNGAseF* (50 U/μL) or RNAse (1:50) but not *O*-glycosidase (4000 U/μL). (D) Cy5-hydrazide labeled sialoglycoRNA signal is reduced in hPAEpC-derived RNA in the presence of the oligosaccharyltransferase inhibitor NGI-1 (20 μM), the α-mannosidase I inhibitor kifunensine (Kif) (15 μM), or the sialyltransferase inhibitor P3-F_Ax_-Neu5Ac (200 μM). Created with Biorender.com

### Lectin labeling reveals α-2,3 and α-2,6 glycosidic-linked sialic acids, and α-2,3 fucose residues in cell surface glycoRNA of alveolar epithelial cells

To elucidate the specific types of glycosidic linkages in the alveolar epithelial glycoRNA, we isolated RNA from the plasma membrane of hPAEpC and labeled it using linkage-specific biotin-conjugated lectins and infrared dye-conjugated streptavidin in a one-pot reaction strategy (Figure 4A). Following incubation, RNA was purified, subjected to denaturing gel electrophoresis, and visualized in-gel, as described above. Alveolar epithelial glycoRNA stained positive with the lectin wheat germ agglutinin (WGA), validating the presence of sialic acid residues (Figure 4B). Positive staining with *Maackia amurensis* lectin (MAA) (Figure 4C) and *Sambucus nigra* lectin (SNA) (Figure 4D) further indicated the presence of both α-2,3 and α-2,6 glycosidically-linked sialic acids within the glycoRNA. Treatment of RNA with neuraminidase or RNAse resulted in the abolition of the sialoglycoRNA signal as detected by any of these three lectins (Figure 4B-D). Additionally, glycoRNA stained positive with *Ulex europaeus* (UEA-1) lectin, suggesting the presence of α-2,3-linked fucose residues, which could be eliminated by treatment with fucosidase (Figure 4E). These findings not only affirm the coexistence of sialic acid and fucose residues, but also the simultaneous presence of α-2,3 and α-2,6 sialic acids within the glycoRNA of alveolar epithelial cells.

**Figure 4.**
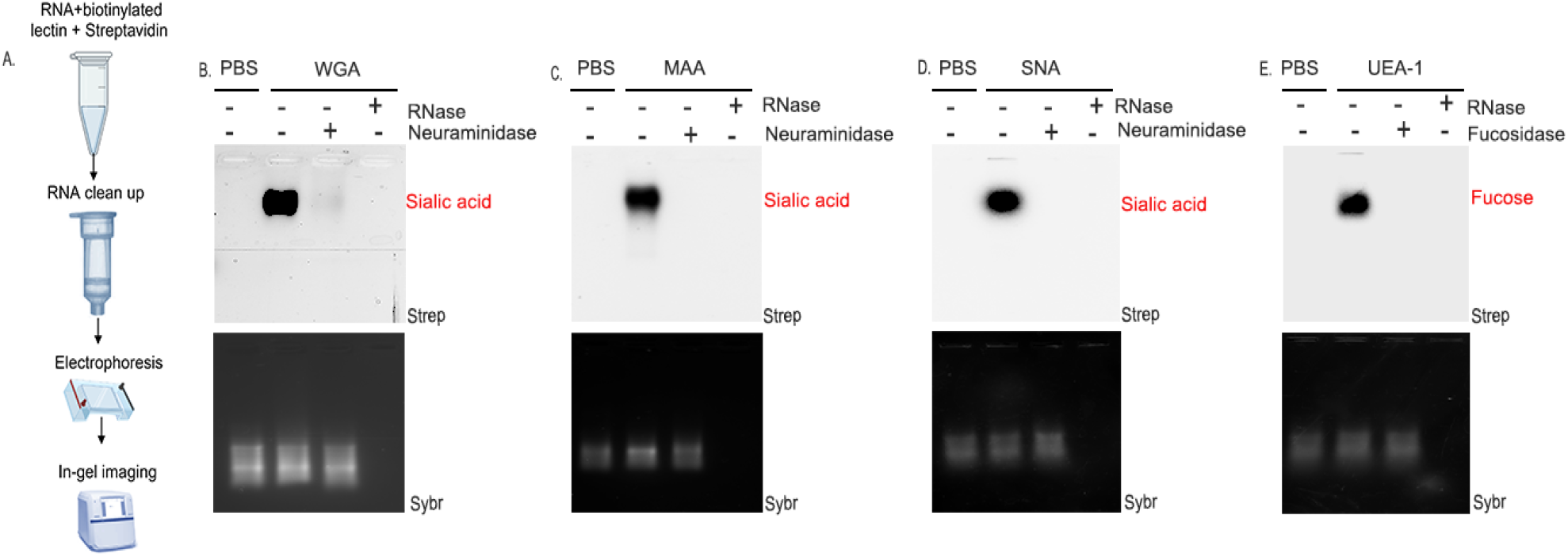
Lectin labeling reveals α-2,3 and α-2,6 glycosidic-linked sialic acids, and α-2,3 fucose residues in cell surface glycoRNA of alveolar epithelial cells. (A) Schematic of cell surface glycoRNA labeling with lectins. RNA was isolated from the plasma membrane of hPAEpC, stained with biotinylated lectins (20 μg/ml) and detected via Streptavidin infrared dye. After purification, 3 μg RNA was loaded onto 1% denaturing agarose gel, electrophoresed for 35 minutes, and imaged on a fluorescent gel scanner. All subsequent gels were treated in this manner. In control experiments, RNA was treated with PBS instead of lectins. RNA was stained with SYBR gold (sybr). (B) In-gel imaging of hpAEpC sialoglycoRNA shows positive staining for wheat germ agglutinin (WGA) that is reduced or lost in the presence of neuraminidase (2 U/μL) or RNAse (1:50). (C) In-gel imaging of hpAEpC sialoglycoRNA shows positive staining for *Maackia amurensis* agglutinin (MAA) that is lost in the presence of neuraminidase (2 U/μL) or RNAse (1:50). (D) In-gel imaging of hpAEpC sialoglycoRNA shows positive staining for *Sambucus nigra* agglutinin (SNA) that is lost in the presence of neuraminidase (2 U/μL) or RNAse (1:50). (E) In-gel imaging of hpAEpC fucoglycoRNA shows positive staining for *Ulex europaeus* agglutinin (UEA-1) that is lost in the presence of fucosidase (0.2 U/μL) or RNAse (1:50). Created with Biorender.com

### Expression of glycoRNAs differ among *in vitro* models of alveolar epithelial cells

To assess the utility of our different fluorescent labeling techniques (namely DBCO-Sulfo-Cy5 following metabolic glycoengineering, Cy5-hydrazide, and streptavidin infrared for biotin-conjugated lectins), we performed a comparative analysis of glycoRNA abundance across different commonly used alveolar epithelial-like cell lines, namely hAELVi, NCI-H441, and A549 cells relative to hPAEpCs. Of these, NCI-H441 cells expressed the highest abundance of sialoglycoRNA, followed by hPAEpC and hAELVi, whereas A549 cells exhibited the lowest levels of both *de novo* formed glycoRNA (as indicated by DBCO-Cy5 labeling, Figure 5A) and total surface sialoglycoRNA (determined by Cy5-hydrazide labeling, Figure 5B). Similarly, *de novo* formed fucoglycoRNA was detected in all cell types with a profound increase in abundance in NCI-H441 cells (Figure 5C). Collectively, these findings reveal distinct cell-type specific variations in glycoRNA abundance across different alveolar epithelial cell lines.

**Figure 5.**
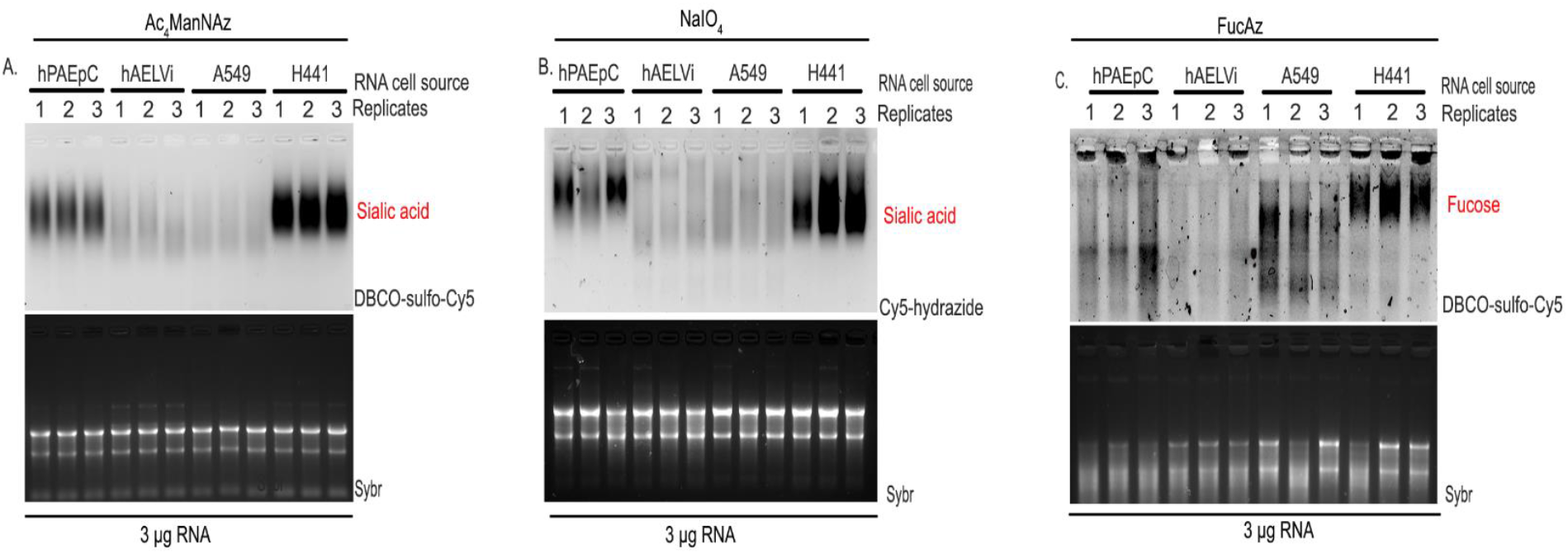
Expression of glycoRNAs differ among *in vitro* models of alveolar epithelial cells. (A) In-gel imaging with DBCO-Sulfo-Cy5 shows differential expression of *de novo* formed sialoglycoRNA in hPAEpC, hAELVi, A549 and NCI-H441 cells pretreated with 100 μM Ac_4_ManNAz for 24 h. (B) In-gel imaging with Cy5-hydrazide shows differential expression of total cell surface sialoglycoRNA in hPAEpC, hAELVi, A549 and NCI-H441 cells subjected to mild periodate oxidation with 1 mM NaIO_4_ for 15 minutes. (C) In-gel imaging with DBCO-Sulfo-Cy5 shows differential expression of *de novo* formed fucoglycoRNA in hPAEpC, hAELVi, A549 and NCI-H441 cells pretreated with 200 μM FucAz for 24 h.

### GlycoRNA contributes to alveolar barrier integrity and viral interaction

To explore potential functional roles of glycoRNA on the surface of the alveolar epithelium, we investigated the impact of enzymatic removal of the glycoRNA on epithelial barrier integrity as well as the interaction with influenza A virus. Treatment with RNAse cocktail depleted cell surface glycoRNA (Fig. 6A) without adverse effects on hPAEpC viability (Figure 6B). Removal of cell surface glycoRNA was associated with a decrease in TEER by approximately 30%, indicating loss of epithelial barrier integrity (Fig. 6C). In parallel, RNAse treatment of hPAEpC reduced the number of detected influenza virus particles (Fig. 6D), suggesting that cell surface glycoRNA contributes to alveolar epithelial barrier function and may act as adhesion molecule for influenza virus attachment.

**Figure 6.**
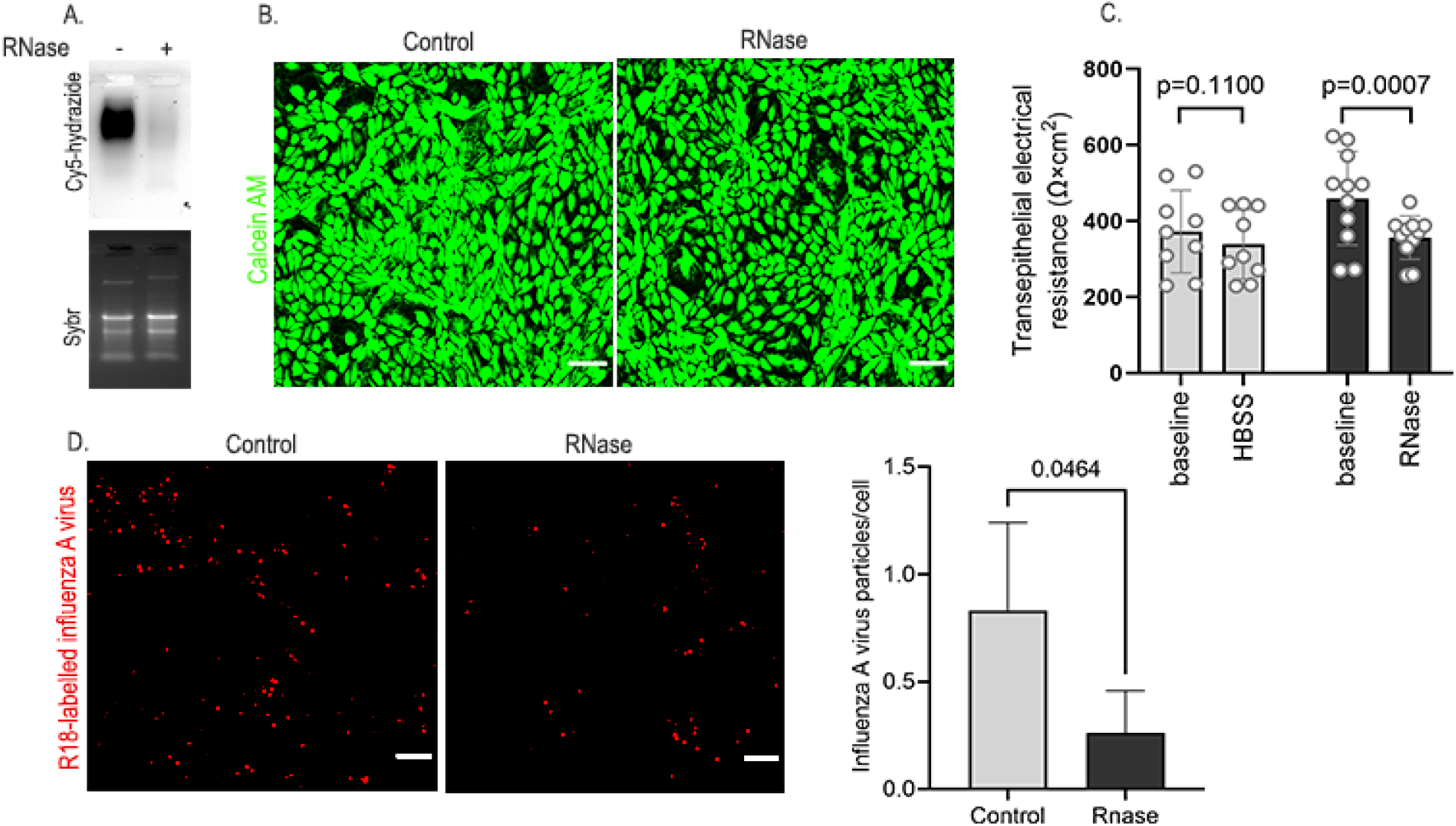
GlycoRNA contributes to alveolar barrier integrity and viral infection. (A) In-gel imaging of sialoglycoRNA with Cy5-hydrazide following mild periodate oxidation shows loss of total cell surface sialoglycoRNA in hPAEpC treated with RNAse cocktail. (B) Staining with calcein AM and propidium iodide show no signs of cell death after RNase treatment relative to HBSS-treated cells. (C) Transepithelial electrical resistance (TEER) is reduced in hPAEpC monolayers treated with RNAse cocktail (1:50). Measurements were taken before and 30 min after treatment. (D) Confocal images and corresponding quantification show reduced abundance of influenza A virus (IAV) particles in hPAEpC treated with RNAse cocktail relative to HBSS control. *p<0.05 vs. control.

## Discussion

Here, we report the first characterization of the expression and function of glycoRNA on the surface of alveolar epithelial cells. To this end, we optimized new techniques to detect glycoRNA on the surface of living cells and in purified RNA by ingel imaging following labeling with different fluorescent dye conjugates. Specifically, we employed DBCO-Sulfo-Cy5 to label newly synthesized glycoRNA following cellular uptake of Ac_4_MAnNAz or FucAz, used Cy5-hydrazide to label total glycoRNA following cell treatment with sodium periodate, and combined infrared-conjugated streptavidin with biotinylated lectins to probe for specific glycan linkages. Leveraging these fluorescent markers allowed for direct visualization of glycoRNA in agarose gels without the need for conventional RNA blotting techniques, and may offer a simple and rapid way to visualize glycoRNA and thus, advance glycoRNA research. Reduced epithelial barrier function and IAV particles reveal novel glycoRNA functions on the alveolar epithelial surface, and identify glycoRNAs as an important component of the alveolar epithelial glycocalyx.

Recent data by Flynn and colleagues identified small RNAs modified with N-glycans on the surface of living cells where they are expected to add to glycoproteins, proteoglycans, glycosaminoglycans and glycolipids in the composition of the glycocalyx ^6^. This discovery has sparked the emergence of new analytical tools for glycoRNA research, and a parallel surge in papers on the expression, composition, and function of glycoRNA on different cells, in different organs, and in various health and disease contexts. Notably among the latter are the recently proposed RNA-glycan linkage via the modified RNA base 3-(3-amino-3-carboxypropyl)uridine (acp3U) as site of attachment of *N*-glycans to glycoRNA ^15^, the identification of an immunomodulatory role of glycoRNAs ^6,7,16^, and their emerging role in tumor malignancy and metastasis ^7^.

In spite of its critical location at the site of gas exchange, surfactant function, cyclic mechanical stress and potential pathogen invasion, the alveolar epithelial glycocalyx has been relatively scarcely studied so far. Previous work by others have shown that loss of alveolar epithelial glycocalyx components such as glycosaminoglycans increases alveolar permeability ^1^ and impairs lung recovery ^17^. Analogously, shedding of glycocalyx components has been found to be associated with alveolar barrier failure in patients with acute respiratory distress syndrome ^1,18^. In parallel, the association of the alveolar epithelial glycocalyx with intraalveolar surfactant ^3^ directly regulates surfactant function and hence, alveolar surface tension ^19^, presumably via direct binding of surfactant proteins to sulfated glycosaminoglycans of the alveolar epithelial glycocalyx ^20^. Expression and function of glycoRNAs on alveolar epithelial cells, and hence, as integral part of the alveolar glycocalyx has so far not been addressed.

In the present work, we demonstrate the expression of glycoRNAs on the cell surface of alveolar epithelial cells. In hPAEpC, cell surface glycoRNA consists of both sialic acids (characterized by both α-2,3 and α-2,6 glycosidic linkages) and fucose (with α-2,3 glycosidic linkage) residues, indicating a complex or hybrid type *N*-glycan composition of the glycoRNA as previously suggested ^6^. A similar compositional pattern was detected in other alveolar epithelial-like cells such as hAELVi, A549, and NCI-H441 cells, with a heightened abundance of glycoRNA in NCI-H441 and reduced abundance in A549 cells relative to hPAEpC. As A549 is an adenocarcinoma cell line, their lower expression of glycoRNAs is in line with a recent report demonstrating an inverse relationship between cell surface glycoRNA expression and breast tumor malignancy ^7^.

Strikingly, the observed differences in glycoRNA abundance across the different cell types in this study parallel their barrier function as reported in our recent study ^12^. Specifically, NCI-H441 cells (with high abundance of glycoRNA) form tight epithelial monolayers, while A549 cells (with low glycoRNA abundance) have a poor barrier function, with hPAEpC ranging in between (in terms of both, barrier integrity and glycoRNA abundance). This observation prompted us to test whether glycoRNA may play a role in epithelial barrier regulation. By TEER measurements following RNAse cocktail-mediated glycoRNA depletion, we detected a decrease in transepithelial electrical resistance by approximately 30%, indicating a relevant contribution of glycoRNAs to epithelial barrier function. This finding is in line with previous data demonstrating the barrier-protective effects of the alveolar epithelial glycocalyx described above. At present, the precise mechanisms by which glycoRNA (as well as other components of the alveolar epithelial glycocalyx) regulate alveolar barrier integrity remain unresolved and should be addressed in future investigations focusing on the interplay between glycoRNA and junctional protein complexes on the one hand, and the role of glycoRNA for the viscosity and permeability of the alveolar lining layer *per se* on the other hand.

Our findings further reveal that depleting cell surface glycoRNA reduces influenza A virus particles in alveolar epithelial cells. This observation is consistent with the well-established role of surface sialoglycans in influenza virus attachment and internalization ^21,22^ but now identifies glycoRNA as a mediator of virus-epithelial interaction. Regulated glycoRNA expression may as such modulate airspace susceptibility to influenza viral infection, and present a putative target for its prevention.

We acknowledge several limitations of our present study. First, we did not conduct detailed molecular analyses on glycoRNA structure. As such, our study does not provide insights into the exact molecular linkage between glycans and RNA in the alveolar epithelial glycocalyx. Our in-gel fluorescent imaging strategies primarily focused on the detection of sialic acids and fucose. Appropriate adaptation of the methods described herein may allow to detect additional sugar residues in glycoRNA. Second, while our lectin labeling suggests the involvement of different glycosyltransferases in the sialylation and/or fucosylation of glycoRNA, the specific enzymes responsible for these modifications in alveolar epithelial cells remain to be determined. Lastly, at the present stage we cannot exclude a possible contribution of non-glycosylated cell surface RNA to the loss of barrier function and viral adhesion following treatment with RNAse.

In conclusion, we report here glycoRNA as a novel component of the alveolar epithelial glycocalyx and a potential regulator of epithelial barrier function and IAV infection. Our findings present numerous avenues for further investigation, e.g. the potential contribution of glycoRNA shedding to changes in alveolar epithelial barrier function in lung injury and pneumonia. As such, measurement of shed glycoRNA in bronchoalveolar lavage fluid could serve as a biomarker for epithelial breakdown and injury. In parallel, the reported role of glycoRNA as ligand for siglecs 11 and 14 on immune cells such as macrophages, monocytes and neutrophils may regulate immune cell dynamics in inflammatory lung diseases. GlycoRNA expressed on the alveolar epithelial surface may also interact with lung collectins (i.e., surfactant protein A and D) analogous to their binding to sulfated glycosaminoglycans recently reported by us ^20^. As such, we consider the identification of functional glycoRNAs on the alveolar surface to bear significant potential to advance and/or revise our understanding of the composition and function of the alveolar epithelial glycocalyx in health and disease.

## Acknowledgements

This study was supported by the Deutsche Forschungsgemeinschaft (DFG, German Research Foundation): project ID 431232613—SFB-1449 subproject B01 to WMK and MO; project ID 437531118—SFB-1470 subproject A04 to WMK; operational grants KU1218/11-1 and 1218/12-1 to WMK; Federal Ministry of Education (BMBF) in the framework of e:Med SYMPATH (01ZX1906A); and the DZHK partner site Berlin to WMK.

We would like to acknowledge Nicole Groenke and Klaus Osterrieder for supporting this work by providing UV-inactivated influenza A viruses.

## Competing interests

The authors declare no competing interests.

## Notes

### Competing Interest Statement

The authors have declared no competing interest.

## References

1. Haeger, S. M. et al. Epithelial heparan sulfate contributes to alveolar barrier function and is shed during lung injury. Am. J. Respir. Cell Mol. Biol. 59, 363–374 (2018).

2. Weidenfeld, S. & Kuebler, W. M. Shedding First Light on the Alveolar Epithelial Glycocalyx. Am. J. Respir. Cell Mol. Biol. 59, 283–284 (2018).

3. Ochs, M. et al. On top of the alveolar epithelium: surfactant and the glycocalyx. Int. J. Mol. Sci. 21, 3075 (2020).

4. Pries, A. R. & Kuebler, W. M. Normal endothelium. Handb. Exp. Pharmacol. 176, 1–40 (2006).

5. Schmidt, E. P., Kuebler, W. M., Lee, W. L. & Downey, G. P. Adhesion Molecules: Master Controllers of the Circulatory System. Compr. Physiol. 6, 945–973 (2016).

6. Flynn, R. A. et al. Small RNAs are modified with N-glycans and displayed on the surface of living cells. Cell 184, 3109–3124.e22 (2021).

7. Ma, Y. et al. Spatial imaging of glycoRNA in single cells with ARPLA. Nat. Biotechnol. 42, 606–616 (2023).

8. Brandt, R. et al. Metabolic Glycoengineering Enables the Ultrastructural Visualization of Sialic Acids in the Glycocalyx of the Alveolar Epithelial Cell Line hAELVi. Front. Bioeng. Botechnology 8, 614357 (2021).

9. Koçak, A. et al. Fluorogenic cell surface glycan labelling with fluorescence molecular rotor dyes and nucleic acid stains. Chem. Commun. 60, 4785–4788 (2024).

10. Park, D. D. et al. Membrane glycomics reveal heterogeneity and quantitative distribution of cell surface sialylation. Chem. Sci. 9, 6271–6285 (2018).

11. Hemberger, H. et al. Rapid and sensitive detection of native glycoRNAs. bioRxiv (2023) doi:10.1101/2023.02.26.530106.

12. Herbst, C. J. et al. Characterization of commercially available human primary alveolar epithelial cells. Am. J. Respir. Cell Mol. Biol. 70, 339–350 (2024).

13. Groenke, N. et al. Mechanism of virus attenuation by codon pair deoptimization. Cell Rep. 31, (2020).

14. Zeng, Y., Ramya, T. N. C., Dirksen, A., Dawson, P. E. & Paulson, J. C. High-efficiency labeling of sialylated glycoproteins on living cells. Nat. Methods 6, 207–209 (2009).

15. Xie, Y. et al. The modified RNA base acp3U is an attachment site for N-glycans in glycoRNA. bioRxiv (2023) doi:10.1101/2023.11.06.565735.

16. Zhang, N. et al. Cell surface RNAs control neutrophil recruitment. Cell 187, 846–860.e17 (2024).

17. LaRivière, W. B. et al. Alveolar heparan sulfate shedding impedes recovery from bleomycin-induced lung injury. Am. J. Physiol. Lung Cell. Mol. Physiol. 318, L1198–L1210 (2020).

18. Sauer, A. et al. Circulating hyaluronic acid signature in CAP and ARDS - the role of pneumolysin in hyaluronic acid shedding. J. Int. Soc. Matrix Biol. 114, 67–83 (2022).

19. Rizzo, A. N. et al. Alveolar epithelial glycocalyx degradation mediates surfactant dysfunction and contributes to acute respiratory distress syndrome. JCI insight 7, (2022).

20. Avcibas, R. et al. Multivalent, calcium-independent binding of surfactant protein A and D to sulfated glycosaminoglycans of the alveolar epithelial glycocalyx. Am. J. Physiol. - Lung Cell. Mol. Physiol. 326, (2024).

21. Zhao, C. & Pu, J. Influence of host sialic acid receptors structure on the host specificity of influenza viruses. Viruses 14, 2141 (2022).

22. Ibricevic, A. et al. Influenza virus receptor specificity and cell tropism in mouse and human airway epithelial cells. J. Virol. 80, 7469–7480 (2006).

